# Age-related decline in metabolite heavy isotope content in rodent organs

**DOI:** 10.1101/724435

**Authors:** Xiyan Li, Michael P. Snyder

**Affiliations:** Department of Genetics, Stanford University School of Medicine, Stanford, CA 94305-5120 USA

## Abstract

Heavy isotopes are discriminated by biological systems due to kinetic isotopic effects at the biochemical/metabolic levels. How these heavy isotopes are enriched or depleted over a long term is unclear, but artificial manipulation of heavy isotope content in various organisms has produced significant impacts on biological functions, suggesting the origin may arise with intrinsic mechanisms for a functional outcome. Our previous study has revealed an age-associated decline in metabolite heavy isotope content (HIC) in the budding yeast, which could be reversed in part by supplementing heavy water, and consequently, also increased yeast lifespans. In the current study, we report a similar age-dependent decline in HIC from three types of mouse tissues: brain, heart, and skeletal muscles. Furthermore, individual tissues exhibited different patterns of HIC change over age, which appeared to match their development and maturation timelines. These results have demonstrated that age-dependent decline in HIC also exists in mammals, which is likely a traceable feature of development and perhaps aging. Thus, we believe that reversing the decline in HIC could have the potential to extend the healthspan of humans.

## INTRODUCTION

Heavy isotopes co-exist as the natural counterparts of common elements (C, H, O, N, S etc.) found in all life forms (Li and Snyder, 2016a). How these heavy isotopes change naturally has not been investigated in a systematic manner until recently, although their effects on biological functions through artificial fortification have been long documented. Despite their relative minor abundance in percentage, these non-radioactive heavy isotopes have profound impacts on biological activities through kinetic isotopic effects, a chemical term describing their ability to affect the dynamics of chemical reactions. Previous studies have focused on manipulating heavy isotope content (HIC) through easily accessible resources, such as supplying heavy water (D2O), in microbes, plants and animals (Katz, 1960; Lewis, 1934). Under high doses of heavy water, slower growth, delayed germination, or slower metabolism was observed, abnormal functions also appeared in cells and organs. However, it seems that some organisms were able to overcome these deleterious effects by adaptation, and potential damages, if any, were reversed when heavy water was withdrawn. Therefore, non-extreme elevation in heavy isotope content does not seem to impose an imminent threat to health.

Living organisms have, besides the heavy hydrogen deuterium, many other types of heavy isotopes. Through a metabolomics study, we have previously examined in amino acids the heavy isotopes for four common organic elements, namely ^13^C, ^2^H, ^15^N and ^24^S, in yeast cells that underwent chronological aging (Li and Snyder, 2016b). We found an overall trend of decline, which occurred when yeast cells get old, and elevating the HIC for hydrogen through heavy water feeding extended yeast lifespan by 84%. Our observation echoed the results of a previous study, which demonstrated that transient heavy water treatment extended lifespan in fruit flies, and more recently, another study showing that ^15^N-enriched food extended lifespan in rotifers (Gorokhova, 2017; Hammel et al., 2013). These studies suggest that HIC elevation may produce a universal pro-longevity effect, but before we could address this directly in mammalian animals, it is still unclear whether an overall HIC decline exists in mammals and for practical reason, whether HIC manipulation would promote longevity in model animals that are closer to human in biology.

In this study, we performed a metabolomics profiling of three vital internal organs of mice at different age points. Our results revealed age-associated amino acid HIC decline in these internal organs, which supports that HIC is an age marker in mammals, and manipulation of HIC may produce similar pro-longevity effects, as those observed in the yeast and fly studies.

## RESULTS

### Study design and measurement of heavy isotope content in metabolites

To examine the physiological metabolism changes that happen at different ages, we profiled the metabolomes of three essential organs, brain, heart, and skeletal muscles, in BALB/C mice of postnatal 1 week, 8 weeks, and 26 weeks (Fig. 1A). These three age points were selected to represent infant, post-puberty, and early adulthood, three important development stages for mammals including humans (Dutta and Sengupta, 2016). No senescent ages were used in this study, because BALB/C mice live an average lifespan of (2 years) years and it was very difficult to obtained aged cohorts from commercial animal organ providers.

**Figure 1.**
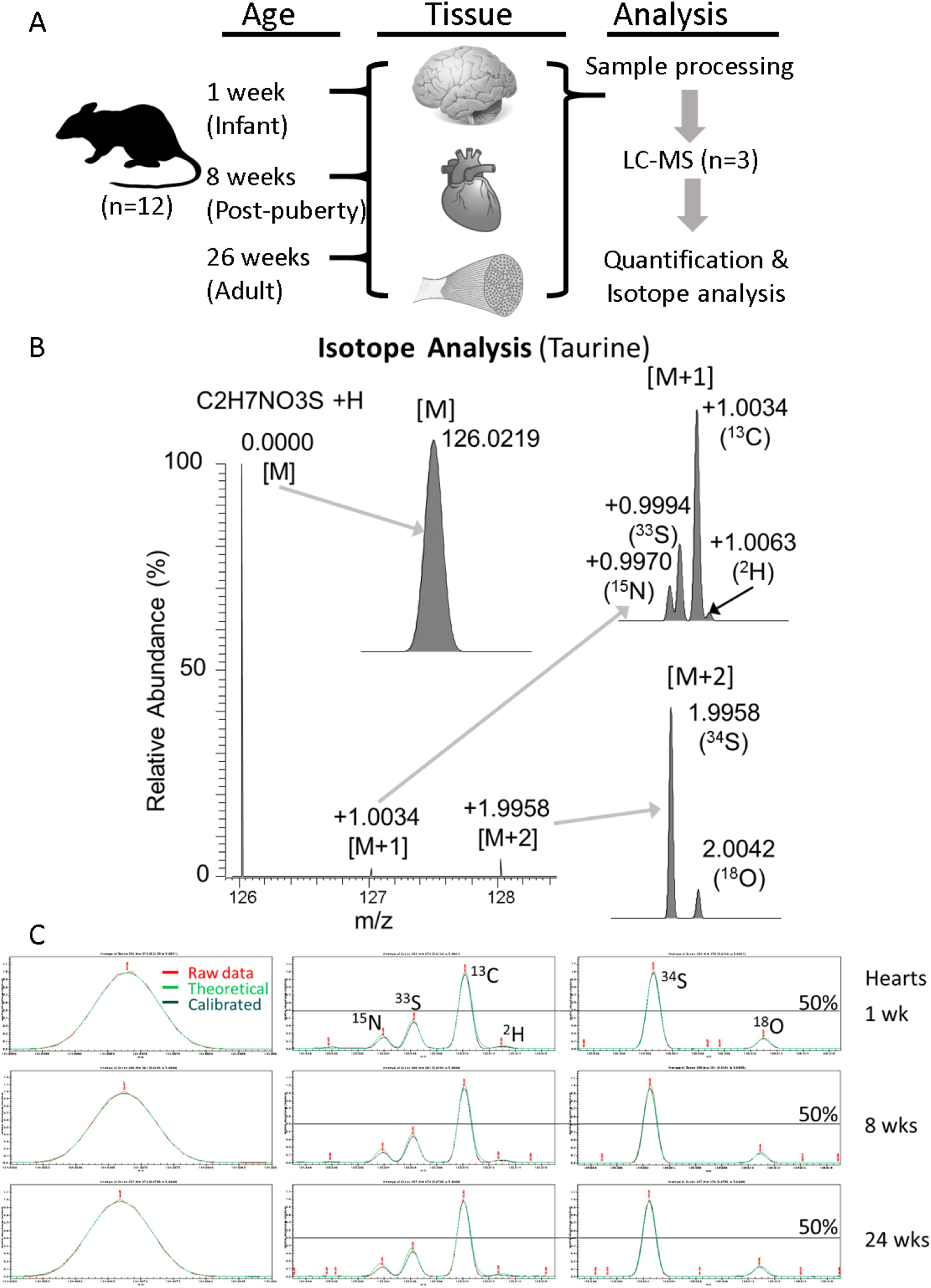
Overview of the study and methodology. **A**. Internal organs and tissues from 12 mice in each age group were analyzed by LCMS (3 technical replicates) for their metabolome and natural heavy isotope content in selected metabolites. **B**. A simulated example demonstrating the natural heavy isotope-containing mass peaks in MS at 100,000 mass resolution for taurine, a sulfur-containing metabolite. **C**. An actual example of the heavy isotope peaks of taurine in mouse hearts analyzed by Massworks in this study. The lines of 50% abundance are labeled as reference. Red represents acquired raw data mass trace. Green represents theoretical mass trace calculated by the software. Blue represents the mass trace calibrated against standards in current experiment settings.

We normalized metabolite extraction against tissue weight to ensure that metabolite measurements across different organs and age groups are directly comparable in this study. Untargeted liquid chromatography-mass spectrometry (LCMS) was performed to simultaneously assess the metabolite composition and heavy isotope contents in metabolites, as previously described (Li and Snyder, 2016b). When operating at a mass resolution of 100,000 and acquired in profile mode, monoisotopic mass peaks can be discerned and measured around the M+1 mass peaks for heavy isotopes ^13^C, ^2^H, ^15^N, and ^33^S, and around the M+2 mass peaks for heavy isotopes ^18^O and ^34^S. One example for the metabolite taurine is shown in Fig. 1B, where predicted relative mass differences for each heavy isotope peaks are indicated.

Our LCMS analyses of mouse hearts indicated that the heavy isotope mass peaks were well separated and thus measurable for quantitative analysis (Fig. 1C). Using 50% abundance as the reference line, we did not observe significant changes of the heavy isotopes in one metabolite taurine in the hearts of 3 different ages. For practical reasons, only ^13^C, ^2^H, ^15^N and ^34^S isotope contents were systematically measured and analyzed in this study.

### Age-related changes in mouse organ metabolome

Through metabolomic analyses using both positive and negative mode acquisitions, we were able to extract 3340 mass features that could be consistently detected in all the samples (3 organs for each of 36 mice, 3 different age groups). Both hypothesis-free model (PCA) and hypothesis-dependent model (OPLSDA) were generated among all organ-age groups to represent the overall clustering patterns (Fig. 2A). Clusters formed clear patterns by organ types in both models, demonstrating the power of our analysis in revealing tissue-specific metabolomics features. Similarly, samples for animals of the same age tend to cluster together, but to a lesser extent, indicating a clear impact of age on metabolome. Our findings are consistent with results of a previous study (Sugimoto et al., 2012), suggesting that individual internal organ possesses unique metabolome signatures, whose chemical compositions likely remain distinctive during development.

**Figure 2.**
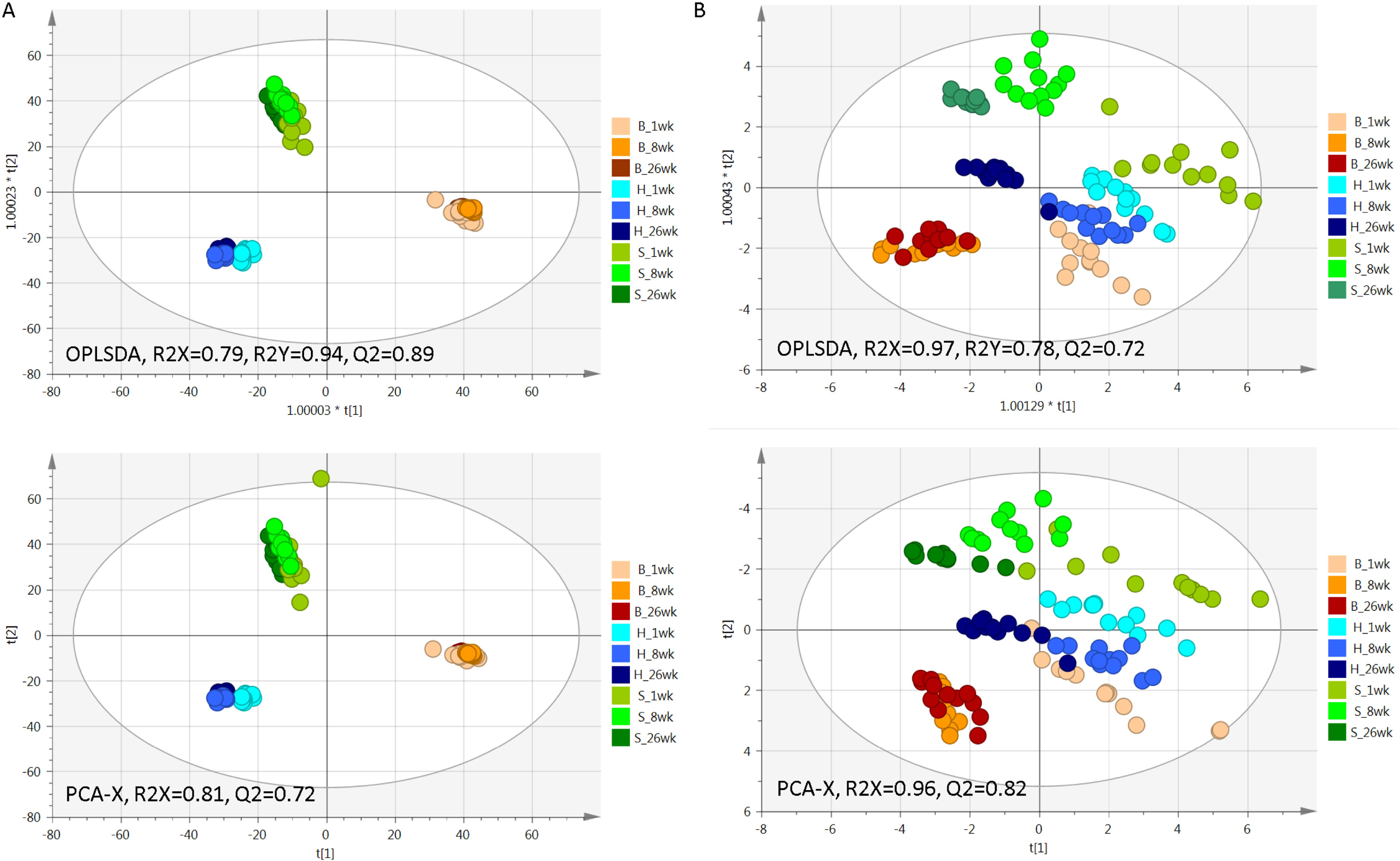
Metabolomics profiles of age-defined mouse organs. Metabolome compositions of mouse brains (B), hearts (H) and skeletal muscles (S) at different ages are clustered using a hypothesis-free PCA-X model or grouping-informed OPLSDA model. Both the whole metabolome (Left, mass feature N=3340) and the 19 confirmed amino acids (Right, metabolite N=19) were used to these clustering plots. Each circle represents an age-specific mouse organ sample. Plots were generated in SIMCA. The white ellipse area represents 95% confidential interval. Model stats are indicated on plots. n = 12 for each age-organ group.

The technical limitation from the instrument, such as mass resolution of 100,000, only allows separation of monoisotopic mass peaks for molecules that are roughly below 250 Dalton, We therefore selected 20 amino acids (our LCMS method did not distinguish leucine or isoleucine) as a representative group and used their isotopic features to generate focused clustering models (Fig. 2B). Besides observing organ-specific clusters using these amino acids, we also noted that different age groups appeared to exhibit similar plot positions for all three organ types, with the youngest groups at the lower right part and the oldest groups at the upper left (Fig. 2B). In addition, different organs exhibited different converging patterns in their amino-acid profiles along ages. The clusters of infant and post-puberty hearts tend to be close to each other, but are at a distance from the cluster of adult hearts. Conversely, the post-puberty and adult brains possessed similar amino-acid profiles, which are significantly different from those present in infant brains. Skeletal muscles showed similar trend as brains, but to a lesser extent.

To further analyze the age-associated metabolomics changes in each organ, we generated O2PLSDA cluster models for the global and focused metabolite groups, using metabolomic features for each organ accordingly. Again, both the global metabolomics (N=3340) and selected metabolomics (N=19, amino acids only) exhibited similar clustering for each organ (Fig. 3A vs. B, respectively), all of each showed a similar transition pattern from young to adult ages. Interestingly, we observed a similar clustering pattern, using both global features and amino-acid profiles; and with only 19 amino acids, we were able to distinguish the profiles of different tissues of different age groups. These results highlight the feasibility and importance to use amino-acid profiles to represent, at least in part, the overall transition of metabolome during mouse development.

**Figure 3.**
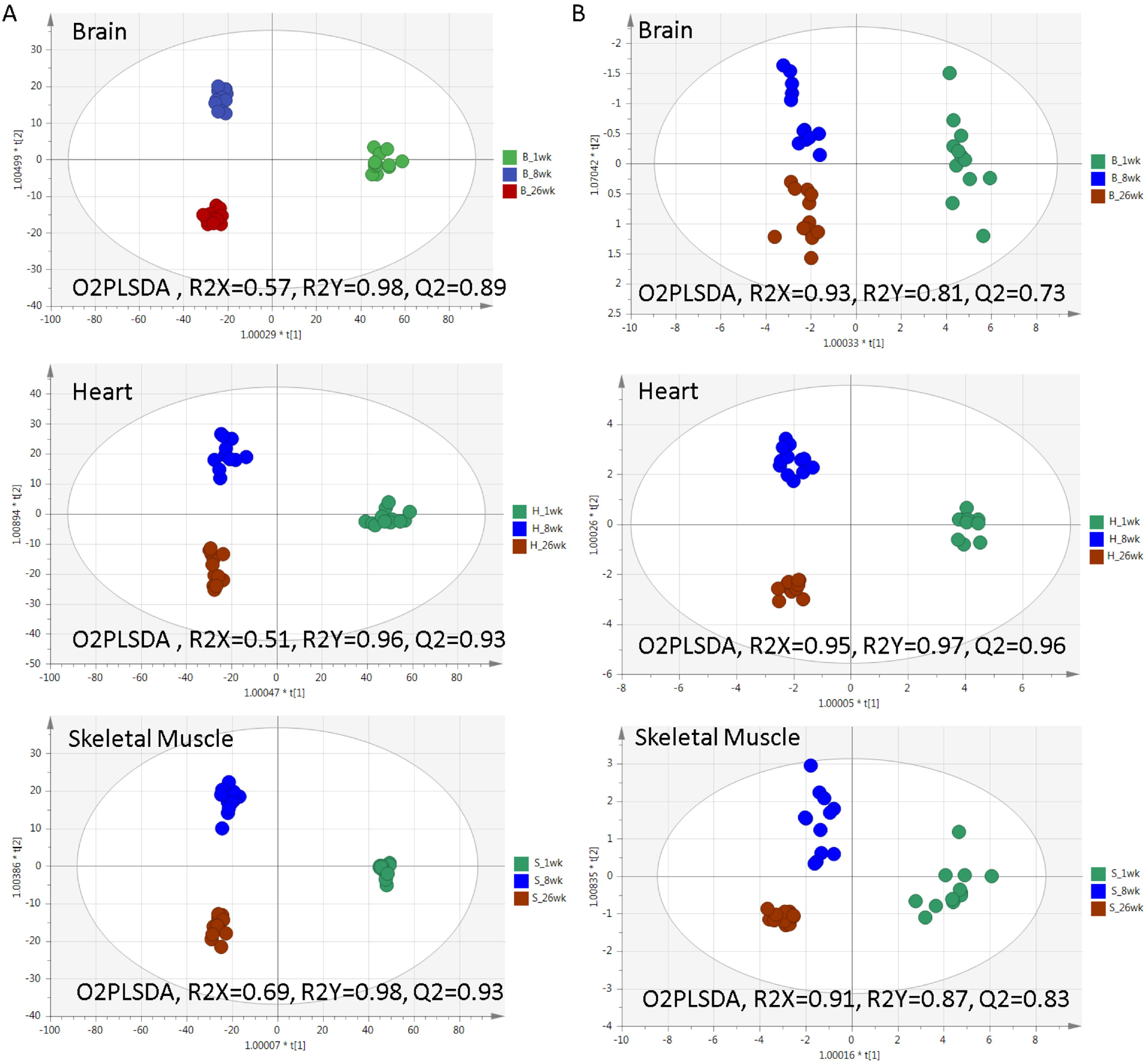
Metabolomics profiles of respective mouse organs by age. Metabolome compositions (left panels: global features; right panels: amino acids only) of individual mouse organs including brains (B), hearts (H) and skeletal muscles (S) at different ages are clustered, using a hypothesis-free PCA-X model or grouping-informed OPLSDA model. See Figure 2 for technical details. n = 12 for each age-organ group.

### Age-related change of individual metabolites associated with redox and urea cycle

To illustrate certain metabolic processes that showed obvious changes among the organ-age groups, we summarized a panel of related metabolites measured in this study (Fig. 4). For example, our results indicated that while NADH increases from infant to adulthood in all three organs, NAD increases drastically only in the heart but not in the brain or skeletal muscles. As a consequence, while NADH/NAD ratio increases by 3-5 fold in the brain and skeletal muscles, it actually drops in adult heart from younger ages, indicating a difference in NAD/NADH metabolism in these organs. Similar age-related decline in the NADH/NAD ratio was also reported in the rat heart previously (Braidy et al., 2011), which was thought to represent a deteriorating capability to maintain intracellular redox status during aging. Our results also showed that this downward trend already manifested at post-puberty age. Drastic increase of NAD in adult heart may also underpin the importance of mitochondria in maintaining heart function and echoes the overall metabolome change (Fig. 2B), because 70% of NAD is mitochondrial in cardiac myocytes, cells that rely on maximal oxidative phosphorylation through mitochondria (Stein and Imai, 2012).

**Figure 4.**
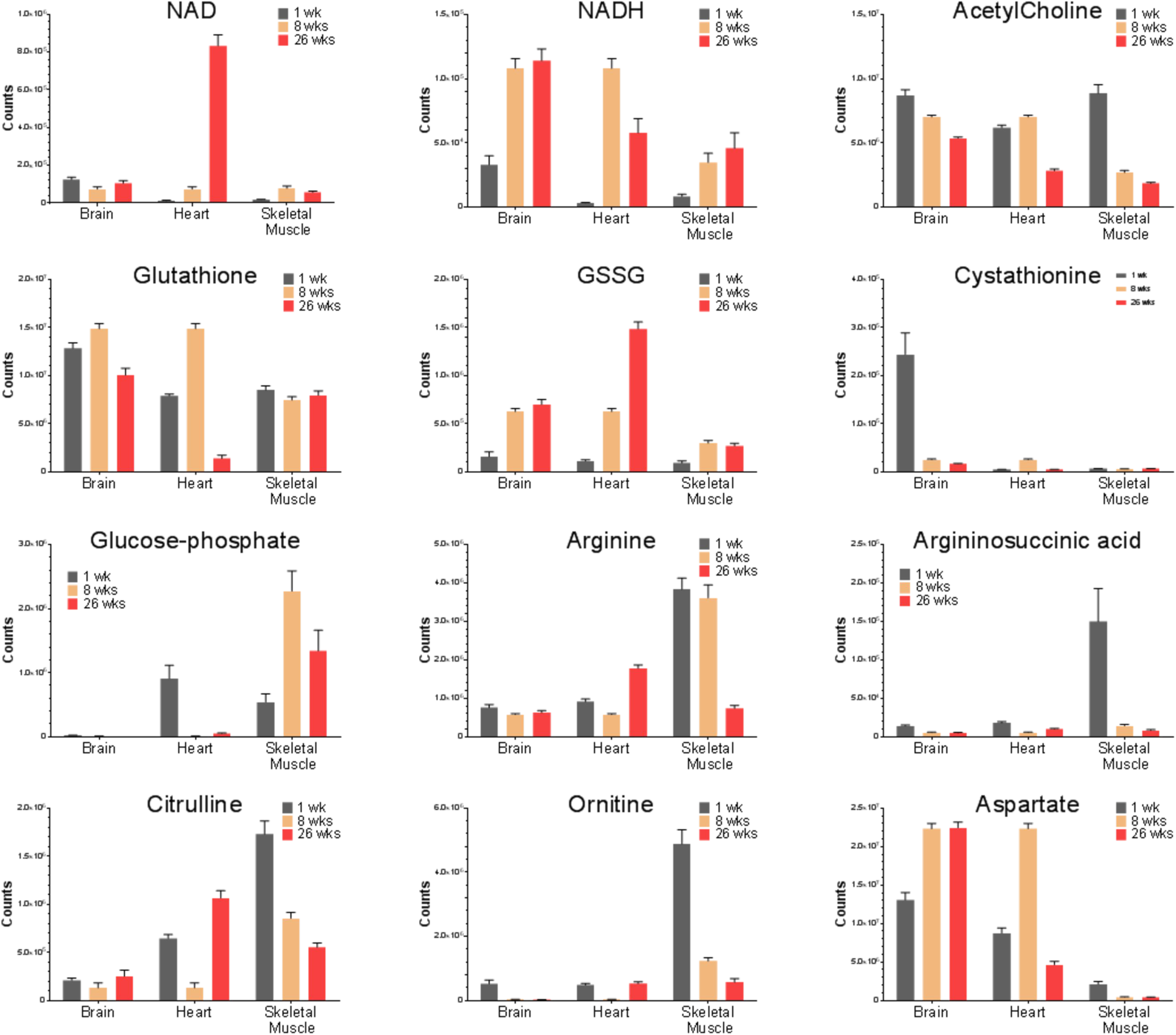
Representative individual metabolite levels in mouse organs at different ages. The metabolite levels were determined by LCMS and plotted across all nine age-organ groups. n = 12. Error bars = SEM. NAD: nicotinamide adenine dinucleotide. NADH: reduced NAD. GSSG: oxidized glutathione. The abundance was normalized by tissue mass. Metabolite names are indicated on the plot. P values from two-tailed t-test (unequal variance, Welch’s correction) were calculated in Graphpad and omitted for clarity.

Similarly, the changes in glutathione and its oxidized form GSSG, another pair of metabolites crucial to redox maintenance, as well as their relative ratios, also mirrored the changes observed for NADH, NAD and NADH/NAD ratios (Fig. 4), although the magnitude of GSSG increase in adult heart was less than that of NAD. Age-related decline in both NADH/NAD and glutathione/GSSG appeared to consistently support the note that deterioration of redox maintenance occurs as early as post-puberty ages in mice.

Interestingly, cystathionine, a metabolite intermediate in methionine to cysteine conversion, showed exceptionally high levels in infant brains (Fig. 4). Previously, it has been shown that human brains contain very high levels of cystathionine, e.g. 68 times as high as in other tissues (Tallan et al., 1958). The cystathionine level is even higher in fetus than in adults (Heinonen, 1976). In comparison, we found the cystathionine level in the brain is 44 times as high as in other tissues. Our results recapitulated the reported changes of cystathionine during the development of mammalian brains and support a characteristic metabolic activity in early brain development that is worth further investigation.

We also found that acetylcholine declined in all three organs with age, with the most drastic change in skeletal muscles (Fig. 4). Since acetylcholine mediates neuron-guided muscle activation and autonomous neuron activities, the observed decline of acetylcholine could be explained by the relative growth of non-acetylcholine requiring tissues in these organs, e.g. more myocytes in muscle enlargement during development.

We also noted that the level of glucose-phosphate increases in skeletal muscles at two later ages, suggesting that the glucose-burning process predominates in mature skeletal muscles, which is contrary to the trend of glucose-phosphate in the heart (Fig. 4). Our method is unable to distinguish between glucose-1-phosphate and glucose-6-phosphate. Nonetheless, the total amount of these two biochemically linked metabolites could also be informative. We believe that the variations suggest an intrinsic difference in energy metabolism between the heart and skeletal muscle. For example, it has been reported that a larger (>10 fold) capacity of gluconeogenesis (anabolism) than glucose phosphorylation (catabolism) occurred in skeletal muscle than in heart (Bass et al., 1969), which is also consistent with the known role of skeletal muscle as a regulator in blood sugar homeostasis.

Strikingly, several metabolites in the urea cycle, namely arginine, arginosuccinic acid, citrulline, ornithine, and aspartate all exhibited an obvious trend of decline in skeletal muscle with age, whereas several metabolites (arginine, citrulline, aspartate) increased in the brain and heart with age (Fig. 4). As urea cycle activity is important to excrete extra nitrogen when amino acids undergo gluconeogenesis, our results suggest an age-related change in urea cycle that may be linked to altered contribution of amino acids to glucose metabolism. Although it is premature to conclude whether it reflects a boost or attenuation of urea cycle at this point, the decline in the anabolic process of urea cycle appears to be consistent with the hypothesis in muscle aging that the anabolic response is gradually lost resulting muscle loss with age (Fry and Rasmussen, 2011).

### Age-related decline in heavy isotope content of amino acids

Since our clustering analysis using only the amino acid profiles could, at least in part, separate different organs of different ages from each other, we wonder whether the common HICs in amino acids could show certain age-related changes, which might also reflect a global change of other metabolites. Taking the advantage of high mass resolution used in our LCMS acquisition, we quantified the common HICs in amino acids as previously described (Li and Snyder, 2016b). As an example, the quantification of total metabolites (total count), as well as the relative abundance of ^13^C, ^15^N and ^34^S isotopic forms of methionine in three organs of three different ages was shown in Fig. 5A. We noticed an obvious age-related decline for methionine containing ^13^C and ^34^S in all three organs, and for ^15^N-methionine in heart and skeletal muscle (note: the level was too low to be determined in the brain). It is unlikely that the age-related decline in the total counts could completely explain the observed decline in the HIC, because the methionine level in heart peaked at 8 week, but steadily declined in HIC, suggesting they are subject to different regulatory mechanisms.

**Figure 5.**
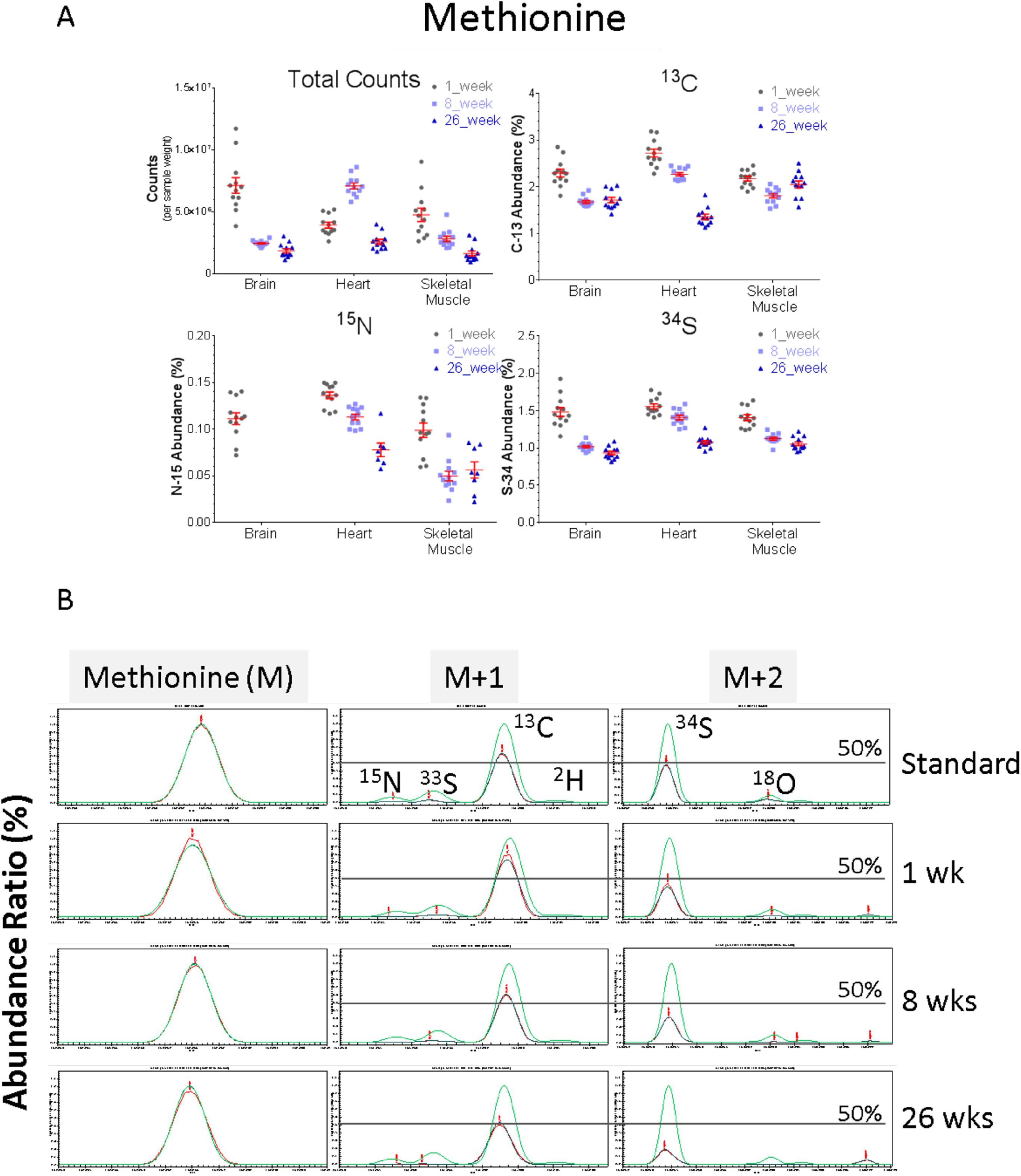
Example heavy isotope content measurement in this study. Top: actual measurement (3 technical replicates per animal replicate) of the heavy isotope-specific metabolite levels, namely ^13^C, ^15^N, and ^34^S, were determined by XCalibur and plotted across all nine age-organ groups. n = 12 for each age-organ group. Error bars = SEM. Bottom: the theoretical and actual heavy isotope-specific metabolite mass curves were determined by MassWorks and plotted across 3 representative heart-age samples. Green trace: theoretical mass curve; Blue: experimentally calibrated mass curve; Red: actual mass curve in the sample. A straight line indicating 50% abundance is added to aid assessment.

To affirm the above quantification results for methionine, we also used MassWorks to assess heavy isotope-containing mass peaks in raw LCMS data after calibration of HIC against respective standard metabolites (Fig. 5B). For methionine again, both raw (red) and calibrated (blue) experimental mass peaks for heavy isotopes dropped below theoretical values (green traces), which is consistent with our previous findings (Li and Snyder, 2016a). Still, both ^13^C and ^34^S peaks showed an obvious decline in hearts with age. These two different methods together validated our observation of an age-related decline in HIC of methionine.

We then applied the same methods to all 20 amino acids. Our method was unable to distinguish leucine and isoleucine, so we combined these two as a single entry. Besides, not all HICs can be reliably quantified for all amino acids due to low counts, such as ^2^H for lysine in the brain. The measurement of total counts and respective heavy isotopes for four amino acids are shown in Fig. 6, including two essential amino acids (valine and lysine) and two non-essential amino acids (glutamine and glutamate).

**Figure 6.**
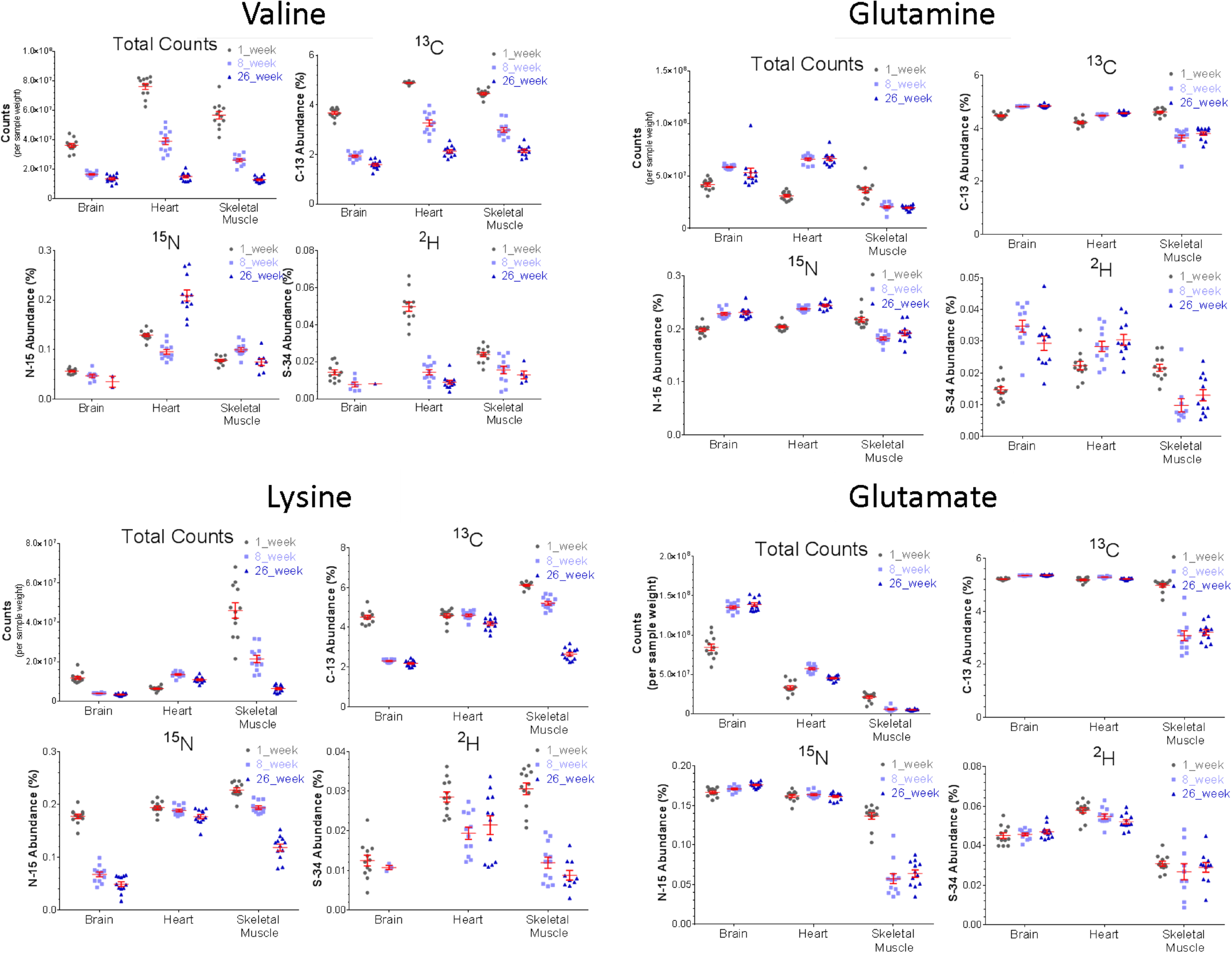
Heavy isotope content measurement in amino acids from age-defined mouse organs. Labels are the same as in Figure 5 top panel. The isotope abundance for some samples was not measurable due to low overall metabolite level. Error bars = SEM.

The total level and ^13^C- and ^2^H-containing forms of valine all declined in all three organs with age, with the most drastic change in heart. In contrast, ^15^N-valine was elevated in the heart (Fig. 6). As a branched chain amino acid (BCAA), valine is actively catabolized in the heart and involved in multiple regulatory mechanisms (Huang et al., 2011). Based on our findings of age-related increase in ^15^N-valine, the nitrogen metabolism of valine, especially transamination, would be a target for further study.

The level of lysine was slightly elevated in the heart but declined in the brain and skeletal muscle with age, whereas all of its HICs declined with age, although to a lesser extent in heart (Fig. 6). The counter-age elevation of lysine in the heart may be associated with its ability to improve contraction in mouse and human hearts (Boldt et al., 2009), possibly as a compensatory mechanism to buffer age-related function decline.

Glutamine and glutamate are among the most abundant and active amino acids in metabolism, thus could best reflect the biochemical discrimination against heavy isotope-containing metabolites. Their levels were both elevated in the brain and heart, but declined in skeletal muscle with age (Fig. 6). However, the HICs in glutamine only, but not in glutamate, changed in the same direction as total metabolite levels in all three organs as age increased. The difference in HIC change between glutamate and glutamine likely reflects their different roles in metabolism: glutamate is a non-essential amino acid, which can be completely depleted from the nutrient supply, whereas glutamine is a conditional amino acid that serves diverse roles in respiration, acid-base balance, carrier of nitrogen, and precursor synthesis of DNA and RNA bases, especially for proliferating cells (Lacey and Wilmore, 1990). In this regard, glutamine phenocopies valine and lysine in their HIC change, probably due to their relatively restricted biosynthesis through metabolism as essential nutrients.

### Overall decline in heavy isotope content of amino acid metabolites

Our study has found that heavy isotope content evidently declined in a group of select small molecule metabolites with age (summarized in Fig. 7). This pattern of HIC decline is consistent with our previous report in yeast cells undergoing aging (Li and Snyder, 2016b), and reflects a pan-kingdom conservation in gradual depletion of HICs with increased age. At present, it is still not clear what biological processes contributed to this decline and what the consequence could be, although some hypotheses, including the role of isotopic effects in metabolism, were discussed (Li and Snyder, 2016a). Moreover, our previous work has suggested that raising cells in a medium rich in heavy isotopes (e.g., deuterium) could substantially extend the replicate lifespan of yeast (Li and Snyder, 2016b). Given its existence in both fungi and mammals, the age-associated heavy isotope content decline may hint on a fundamental principle in the biochemical reaction system known as metabolism during the processes of development and aging.

**Figure 7.**
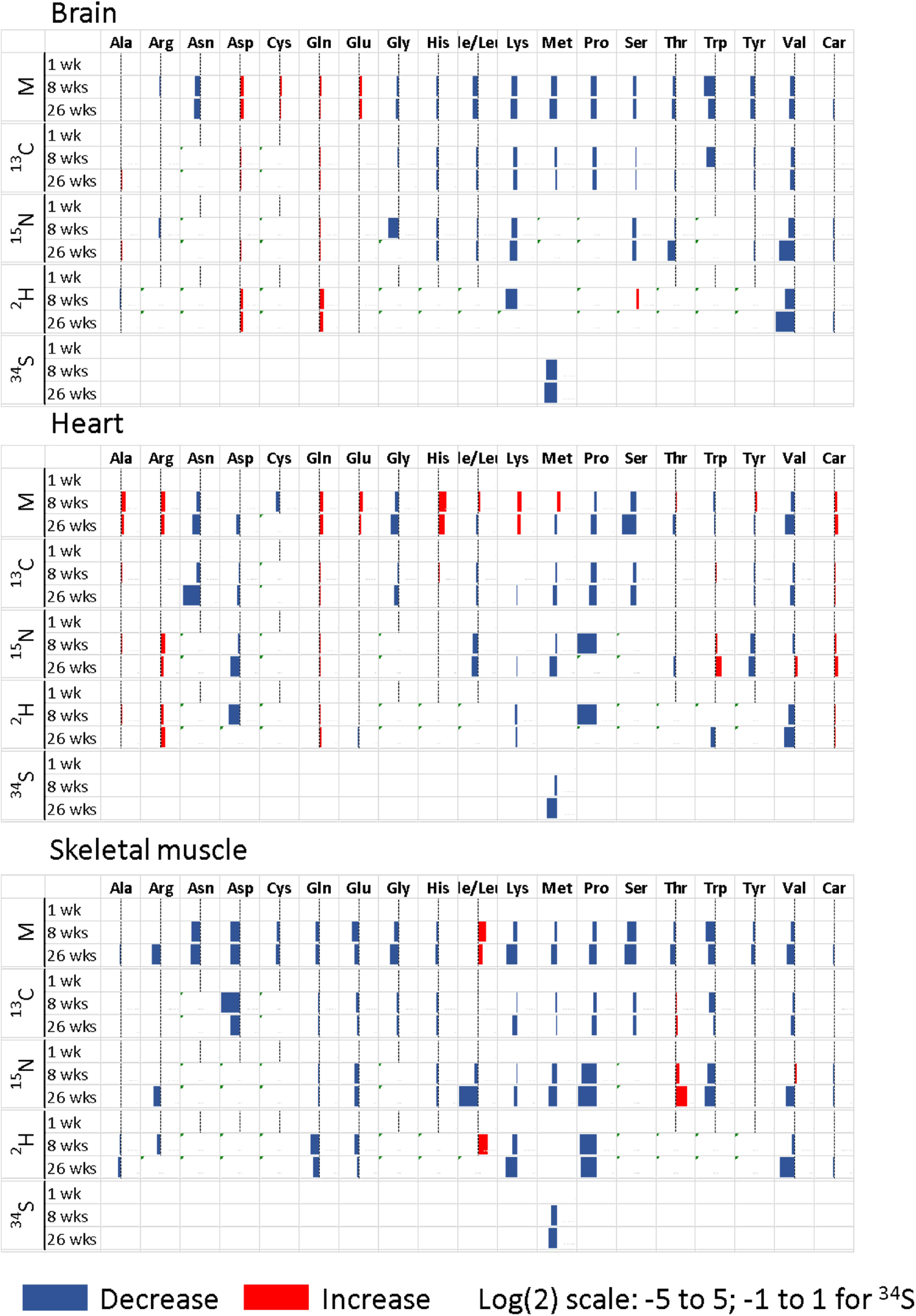
Summary of heavy isotope content in amino acids from mouse organs of different ages. Bar plots of log2 scale of the average fold change (n=12) of the heavy isotope content of each of 19 amino acids and carnitine at age 8 weeks, 26 weeks versus age 1 week. Car, carnitine. .

## DISCUSSION

Heavy isotope content is both a measurable and actionable feature with respect to aging. In essence, the current study in mammalian tissues recapitulated the overall HIC decline in amino-acid metabolites we have reported in the yeast (Li and Snyder, 2016b), disclosing a fundamental chemical basis that is conserved between kingdoms. Although no mice at senior age (more than 1 years old) were used in this study, the age-associated decline in HIC is even more intriguing because it may suggest a universal chemical process that spans development and aging. We have discussed similar studies and possible theories for HIC underneath aging elsewhere (Li and Snyder, 2016a). We have postulated that HIC serves as an aging marker and demonstrated that supplementation of deuterium extended yeast lifespan by more than 80% (Li and Snyder, 2016b), which may be expanded to development from the observation in this study. It is still too early to postulate whether and what causal relationship exists, yet at least, we believe that HIC represents a traceable footprint of metabolism, which may see its application in assessing the function states (or health) of the whole body, individual organs, and maybe, specific types of cells.

Heavy isotope content can be elevated with tolerable long-term health risks. Increasing HIC in mammalians through heavy water feeding have caused intoxication effects at a high dosage (4-5 liters for human), but heavy water itself “is never toxic to any high degree” (Lewis, 1934). Long-term feeding of heavy water elevated the plasma deuterium level to 25% in small dogs and caused low sugar metabolism rate, anemia and heart damage. But the damages were “reversible” after the dogs were restored to drinking normal water for a few months (Katz, 1960). These studies have elicited a high HIC condition (25% deuterium) in vivo, which seemed to be well tolerable by the animals and the signs of health risks also appeared to be reversible. Even though the high cost could be prohibitive for testing HIC manipulation in aging experiments, further testing on potential pro-longevity effects by HIC elevation in experimental mammals is still tempting.

Despite the overall age-associated HIC decline, each of the three organs displayed a distinct pattern over timing (Figs. 2 and 3), which coincides with their respective development program and could be due to the traces left from their respective trajectory of metabolism. Our observation in current study suggests that the heart reach its mature stage at a later age than the brain and skeletal muscles, which is consistent with several previous reports showing that mouse heart reached adult volume at postnatal 12 weeks (Leu et al., 2001), whereas the brain started its peak growth at 7 weeks (Semple et al., 2013) and skeletal muscle fibers established adult state at 3 weeks postnatally (White et al., 2010).

Due to technical limitation in the mass spectrometry technology to handle the mass peak intensity in relative to the chemical noises, the net decline of HIC in metabolites may be less than reported when the metabolite level also declines. However, given our findings that HIC decline was also observed in some metabolites that increase with ages, such as methionine, while no HIC decline was seen for glutamate, whose levels increased with ages, our assessment of the trend in HIC change is valid. Further technical development will be needed for more accurate measurement of HIC in small-molecule metabolites.

## ACKNOWLEDGEMENT

This work was supported by NIH grant (5R01GM06248012) and California Institute for Regenerative Medicine grant (RB4-06087) to MS.

The authors would like to thank Dr. Peichuan Zhang for critical comments of the manuscript.

## MATERIALS AND METHODS

### Animal tissues

Mouse tissues were provided by Charles River Laboratories (Wilmington, MA). The C57BL/6 mice were fed with the standard 5L79 diet ad libitum until sacrificed at specified ages. The tissues were provided by Charles River. Fresh tissues were flash frozen in liquid nitrogen and stored at −80C before processing.

### Sample preparation

Frozen tissues were mixed with calculated volume (500-1200 μL, at the ratio of 4:1) of 80% methanol (MS-grade) and lysed on a FastPrep-24 Instrument (MP Biomedicals, Burlingame, CA), using the lysing matrix D and suggested settings (speed 6 and 4 × 30 s pulses) with 1 min interval on ice. The lysate was incubated at −20°C for 15 min before being centrifuged at 22,000 rpm at 4°C for 15 min. The supernatant was transferred to a fresh tube and centrifuged again at 22,000 rpm at 4°C for 15 min. Each sample was further diluted in 80% methanol (MS-grade) to bring the final concentration to 20 μL/mg tissues for LCMS analyses.

### LCMS

Untargeted LCMS analysis was performed as previously reported, using a platform that consists of Waters UPLC-coupled Exactive Orbitrap Mass Spectrometer (Thermo), equipped with an OPD2 HP-4B column (4.6 × 50 mm) with a 4.6 ×10mm guard column maintained at 45°C (Shodex, Showa Denko, Tokyo, Japan) (Li and Snyder, 2016b). The temperature in the sample chamber was maintained at 4°C. The binary mobile phase solvents were: A, 10 mM ammonium acetate, in 10:90 acetonitrile water; B, 10 mM ammonium acetate, in 90:10 acetonitrile:water. Solvents for positive mode acquisition were modified with 10 mM acetic acid (pH 4.75), and for negative mode acquisition, with 10 mM ammonium hydroxide (pH 7.25). The flow rate was 0.1 ml/min and the gradients were: 0-15 min, 99% A; 15–18 min, 99% to 1% A; 18-24 min, 1% A; 24–25 min, 1% to 99% A; 25–30 min, 99% A. The MS acquisition was performed in profile mode with an electrospray infusion (ESI) probe, operating with capillary temperature at 275 °C, sheath gas at 40 units, spray voltage at 3.5 kV for positive mode and 3.0 kV for negative mode, capillary voltage at 30V, tube lens voltage at 120V and skimmer voltage at 20V. The mass scanning was set at 100,000 mass resolution, high dynamic range for AGC target, 500 ms as the Maximum Inject Time and 70-1,000 m/z as the scan range. All data files were generated from a multi-day single batch acquisition without interruption.

### Data analysis

For metabolomics analysis, the raw LCMS data files were centroided with PAVA program and converted to mzXML format by an in-house R script as previously described (Li and Snyder, 2016b). Mass features were extracted with XCMS v1.30.3.45. Tentative metabolite identification was achieved by manual search against the Metlin metabolite database as previously described (Li and Snyder, 2016b).

### Heavy isotope content measurement

Heavy isotope-content in metabolites were quantified directly in Xcalibur Qual Browser v2.2 (Thermo) by using Peak Detection algorithm Genesis and universally-set accurate mass ranges of heavy isotope-metabolites (mass filters attached). The identities of all reported metabolites with heavy isotope measurement were also confirmed by matching with standard compounds.

Alternatively, the heavy isotope mass curves in raw data were also compared with calibrated and theoretical mass curves with specialized metabolite isotope analysis algorithm in MassWorks v5.0 (Cerno Bioscience, Norwalk, CT).

### Data Plot

The scored mass features were clustered with SIMCA v13.03 (Umetric, Malmö, Sweden).

